# No modulatory effect of rhythmic visual alpha/gamma stimulation in a bistable motion illusion

**DOI:** 10.1101/2020.06.07.139287

**Authors:** Michel J. Wälti, Marc Bächinger, Nicole Wenderoth

**Affiliations:** Neural Control of Movement Lab, Department of Health Sciences and Technology, ETH Zurich, Switzerland; Cognition, Perception and Behaviour in Urban Environments, Future Cities Laboratory, Singapore-ETH Centre, Singapore; Neuroscience Center Zurich (ZNZ), University and ETH Zurich, Zurich, Switzerland

**Author notes:** **Correspondence:** Michel J. Wälti.

**Keywords:** perception, bistable motion illusion, stroboscopic alternative motion, motion quartet, steady-state evoked potentials, entrainment, neural oscillations

## Abstract

The ‘motion quartet’, also called stroboscopic alternative motion (SAM), is a bistable apparent motion paradigm where dots are perceived as moving either horizontally or vertically. These two percepts are known to switch randomly depending on both alpha (8 – 12 Hz) and gamma (30 – 80 Hz) oscillations within and between hemispheres, respectively. However, whether these oscillations play a causal role in triggering perceptual switches remains unknown.

Here, we aimed to entrain brain oscillations in the visual system by presenting flickering stimuli. Two LEDs were flickering either at 10 Hz, 40 Hz or corresponding jittered stimulation frequencies on the left and right sides of the visual field during the presentation of a motion quartet. In our control condition (no stimulation), we show that interhemispheric gamma connectivity (spectral coherence) between the visual cortices of both hemispheres is important for sensory feature integration. On the other hand, alpha power did not predict perceptual switches for the motion quartet. Rhythmic visual stimulation with 10 Hz and 40 Hz evoked resonance-like neural responses but did not alter visual perception.

Thus, although sensory stimuli seem to entrain ongoing brain rhythms, the effect on human behavior differs from the principles observed for endogenous brain oscillations.

## Introduction

Conscious human perception is assumed to be established by the synchronization of oscillatory neuronal brain activity both within local areas and across several remote cortical regions (Engel, Fries, & Singer, 2001). In the visual system, the perception of a consistent single visual field is thought to be facilitated by interhemispheric synchronization between early visual areas of both hemispheres (Bressler, Coppola, & Nakamura, 1993; Gazzaniga, 2000; Mima, Oluwatimilehin, Hiraoka, & Hallett, 2001; Srinivasan, Russell, Edelman, & Tononi, 1999). For certain stimuli requiring bilateral visual perception, visual awareness can alternate between two different percepts, even if the physical properties of the stimulus are unchanged. Examples of such ambiguous illusions are the Necker Cube, structure-from-motion (SFM), the spinning wheel illusion or stroboscopic alternative motion (SAM; also: motion quartet), whose underlying neuronal mechanisms of bistable perception have been extensively studied (for reviews, see Leopold & Logothetis, 1999; Sterzer, Kleinschmidt, & Rees, 2009). The bistable perception in SAM is somewhat unique because it either requires (i) the unilateral processing of a perceived motion separately for each of the visual hemifields when vertical movements are perceived, or (ii) an interaction between both hemispheres when the motion of the visual stimulus crosses the retinal midline so that a horizontal movement is perceived (Genc, Bergmann, Singer, & Kohler, 2011; Ramachandran & Anstis, 1983; Rose & Büchel, 2005).

Electrophysiological studies in humans identified two main frequency bands that relate to such perceptual ambiguities: alpha (8 – 12 Hz) and gamma (30 – 80 Hz). While alpha band activity is thought to regulate neuronal network excitability through functional inhibition or destabilization of the actual percept (Flevaris, Martinez, & Hillyard, 2013; Foxe & Snyder, 2011; Jensen & Mazaheri, 2010; Lange, Keil, Schnitzler, van Dijk, & Weisz, 2014), gamma band activity has been proposed to be related to interhemispheric communication (Hipp, Engel, & Siegel, 2011; Rose & Büchel, 2005). It has been shown that a decrease of inhibitory alpha power is correlated with an increased probability of a perceptual switch (Helfrich et al., 2016; Isoglu-Alkaç et al., 2000; Isoglu-Alkac & Struber, 2006; Piantoni, Romeijn, Gomez-Herrero, Van Der Werf, & Van Someren, 2017; Sangiuliano Intra et al., 2018; Strüber & Herrmann, 2002) and that synchronized gamma activity (also: coherence) increases during interhemispheric communication (Bressler et al., 1993; Doesburg, Kitajo, & Ward, 2005; Mima et al., 2001; Srinivasan et al., 1999; Strüber, Başar-Eroglu, Hoff, & Stadler, 2000).

In recent years, methods for entrainment of brain oscillations have been proposed to alter perception by modulating gamma (Cabral-Calderin, Schmidt-Samoa, & Wilke, 2015; Strüber, Rach, Trautmann-Lengsfeld, Engel, & Herrmann, 2014) and alpha oscillations (Helfrich, Schneider, et al., 2014). While various non-invasive methods have been proposed to be capable of frequency-specific modulation of brain rhythms (for a review, see Thut, Schyns, & Gross, 2011), transcranial alternating current stimulation (tACS) has been predominantly used to alter the perception of ambiguous stimuli (for a review, see Herrmann, Rach, Neuling, & Strüber, 2013). Using tACS, Strüber et al. (2014) found that externally desynchronizing gamma oscillations between occipital hemispheres impaired interhemispheric integration of perceived motion (Strüber et al., 2014). Similar results were described by Helfrich et al. (2014) who modulated interhemispheric connectivity with gamma band tACS in a phase-specific way. While in-phase stimulation was accompanied by enhanced synchronization, anti-phase stimulation impaired synchronization. In addition to their gamma-coupling effect, they revealed a decrease in alpha power in response to gamma stimulation (Helfrich, Knepper, et al., 2014).

An alternative form of frequency-specific modulation of brain rhythms can be achieved with rhythmic sensory stimulation, which induces steady-state evoked potentials (SSEPs) following the temporal frequency of the driving stimulus (Regan, 1977). SSEPs can be measured with electroencephalography (EEG) and have been documented in the visual (steady-state visually evoked potentials, SSVEPs), the auditory (auditory steady-state responses, ASSRs) and the somatosensory (steady-state somatosensory evoked potentials, SSSEPs) systems (for a review, see Vialatte, Maurice, Dauwels, & Cichocki, 2010). Similar to other methods of neural entrainment, the underlying mechanism of SSEPs is still debated (see Zoefel, Ten Oever, & Sack, 2018). While some studies report evidence for entrained ongoing brain oscillations by showing an interaction between stimulation and endogenous activity (Notbohm, Kurths, & Herrmann, 2016; Schwab et al., 2006; Wälti, Bächinger, Ruddy, & Wenderoth, 2019), others found responses to be only regular repetitions of evoked neural potentials (Capilla, Pazo-Alvarez, Darriba, Campo, & Gross, 2011; Keitel, Quigley, & Ruhnau, 2014). In contrast to overlaying evoked potentials, entrainment modulates ongoing brain oscillations. However, to convincingly demonstrate that SSEPs actually represent neural entrainment, a dependency on endogenous oscillations might not be sufficient to fundamentally distinguish entrainment from a rhythmic series of evoked potentials (Haegens & Zion Golumbic, 2018). With regard to the ongoing debate on whether neural entrainment occurs as response to rhythmic sensory stimulation, establishing whether changing brain oscillations with SSVEPs modulates behavior in a predictable manner seems to be an important outcome measure (Haegens & Zion Golumbic, 2018).

The goal of the present study was to test whether oscillatory brain activity can selectively be modulated with flickering visual stimuli and alter the perception of a bistable motion illusion. In line with previous research, we focused on alpha (10 Hz) and gamma (40 Hz) stimulation frequencies, applied to both visual hemifields.

## Materials and methods

### Participants

A total of 34 healthy participants with normal or corrected-to-normal vision were recruited and participated in the experiment. 1 participant was excluded from the analysis due to technical problems during the EEG recording and 4 others were not able to perceive a change in motion of the dots during baseline testing of the SAM. This resulted in a final sample of 29 participants (female: 14; age: M ± SD = 24.5 ± 3.4). Participants were naïve to the hypothesis of the study and that the bistability of the SAM was merely a perceptual phenomenon and not a difference in stimuli presentation. This was important because it has been shown that perceptual switches of a bistable stimulus can to some extent be under the voluntary control of the participant (Kohler, Haddad, Singer, & Muckli, 2008; Sangiuliano Intra et al., 2018).

### Bistable motion illusion: SAM

In the bistable motion paradigm used in this study, the perception of apparent motion was induced by flashing two dots that simultaneously appeared in diagonally opposite corners of a virtual rectangle with a constant vertical distance and variable horizontal distance. Dots were alternately switched on and off between the corners of both diagonal axes. This causes an apparent motion of the dots that can be perceived either along the vertical or the horizontal axis.

### Determination of parity ratios

Previous studies presenting SAM over longer time periods (until a perceptual switch was perceived) have shown that the ratio between vertical and horizontal distance that leads to equal durations of vertical and horizontal motion perception can vary strongly between observers (Chaudhuri & Glaser, 2009; Kohler et al., 2008). To determine individual ratios at which vertical and horizontal motion perception were equally likely (parity ratio, PR), we presented SAM before the start of each block of the main experiment at 10 vertical-horizontal distance ratios. The moving dots were presented for 3 cycles and participants were asked to indicate with a key press whether they perceived horizontal or vertical movement of the dots. The 10 ratios were presented in random order and the procedure was repeated in total 4 times. Averaged across the 4 runs, this resulted in a ratio of vertical motion perception for each of the 10 ratios. We calculated individual parity ratios by extracting the value at 50 % likelihood of perceiving vertical or horizontal movement along the resulting sigmoid distribution of the data points (see Figure 1 B).

**Figure 1.**
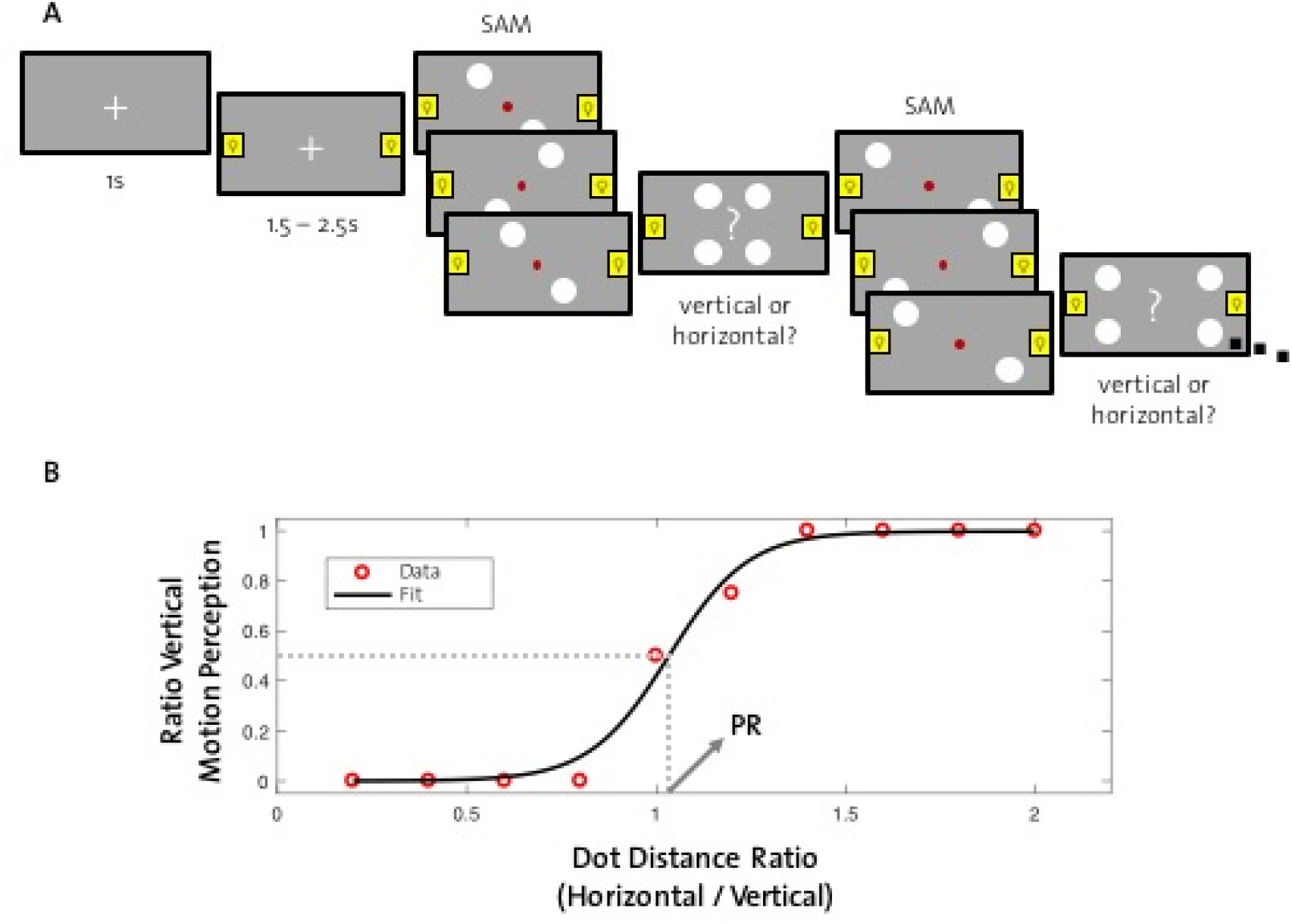
Design of the main experiment and determination of individual parity ratios. A: A trial is depicted with the first two (out of 9) different horizontal distances of the presented dots during SAM. Each SAM is followed by a decision screen, where the subject indicates whether horizontal or vertical movement of SAM was perceived. Horizontal distances are customized to participants parity ratio and adjusted for each block. B: Prior to each block, parity ratio was determined for each subject by presenting SAM at several different dot distance ratios (horizontal distance divided by vertical distance), which generates a sigmoid distribution of data points. Depicted are data from an example subject (red circles) and the corresponding fit of a logistic function (black line). The ratio at which vertical motion perception is at 50 % determines the subject’s parity ratio (PR, here: 1.03).

### Bilateral visual stimulation

Bilateral stimulation of the visual system was caused by means of two white light emitting diodes (LEDs) with a diameter of approximately 2 mm that were mounted on the left and right side of the computer screen. Left and right LEDs flickered in-phase with either 10 Hz, jittered around 10 Hz, 40 Hz, or jittered around 40 Hz frequencies. Stimulation intensity was kept constant across all stimulation conditions.

### Main experiment

Figure 1 A depicts the procedure of the main experiment. For the SAM, varying distances between the dots along the horizontal axis were used, while distance along the vertical axis was held constant. 9 horizontal distances derived from individual parity ratio (PR) were used (PR, PR ± 0.05 x PR, PR ± 0.2 x PR, PR ± 0.4 x PR, PR ± 0.6 x PR). Presentation of SAM was followed by a screen depicting a question mark and all 4 statically presented dots (decision screens). Participants were instructed to indicate whether they had perceived vertical or horizontal motion of the dots by a key press. The experiment was conducted in 4 blocks, each consisting of 2 runs (including all 9 distance ratios presented in random order) per condition (10 Hz, 10 Hz jittered, 40 Hz, 40 Hz jittered, control), presented in pseudo-randomized order. Each of the blocks was preceded by the determination of the individual parity ratio as described before.

### EEG data acquisition

EEG data were acquired at 1000 Hz using a 64-channel Hydrocel Geodesic EEG System (Electrical Geodesic Inc., USA), referenced to Cz (vertex), with an online Notch filter (50 Hz) and high-pass filter at 0.3 Hz. Impedances were kept below 50 kΩ.

### Preprocessing of EEG data

EEG data were preprocessed and analyzed offline. Data were cleaned (detection and interpolation of bad electrodes), band-pass filtered (0.5 – 85 Hz) and further processed using independent component analysis (ICA). Artifact components (ICs) were automatically detected and removed from the data using a custom toolbox (see Liu, Ganzetti, Wenderoth, & Mantini, 2018).

### Behavioral data analysis

In order to determine the rate of both directions of motion perception, we counted the number of vertical percepts and divided this value by the total amount of perceptual choices across trials (perception rate of vertical motion). Next, we subtracted the rate of vertical perception in the control condition from the rates in each of the stimulation conditions (Δ perception rate of vertical motion). A one-factor repeated measures ANOVA analysis was then used to determine differences across stimulation conditions.

A similar approach was followed to determine whether the number of perceptual switches was modulated by our stimulation conditions. For each trial (except the first) we determined whether the response (vertical or horizontal) was equal or switched in comparison to the previous trial. For each condition, we counted the number of perceptual switches across trials (maximal number of switches = 64). The number of perceptual switches in the control condition was then subtracted from the number of perceptual switches in each experimental condition (Δ perceptual switches). Again, a one-factor repeated measures ANOVA was used for statistical analysis.

### EEG data analysis

First, we tested the hypothesized effects of gamma connectivity and alpha power. To do so, data from the control condition were epoched around the onset of each SAM presentation (−500 ms to + 1000 ms) and analyzed.

For gamma connectivity, spectral coherence was assessed in averaged data from 5 occipital electrodes from each hemisphere. Spectral coherence is a measure of phase-based connectivity with phase values being weighted by power values (see Cohen, 2014). Coherence was analyzed across the whole epoch for each trial and then averaged within the low gamma frequency band (36 – 44 Hz). Next, individual trials were divided in two equally-sized groups based on gamma connectivity (low and high coherence). For each subject, we determined the rate of vertical motion perception in both of the groups, which were then statistically compared using a paired-samples t-test.

Alpha power in the control condition was calculated as averaged power across 500 ms preceding the onset of each SAM. For power extraction, a family of complex Morlet wavelets spanning 6 – 14 Hz (in 20 steps) was used with wavelet cycles increasing logarithmically between 4 and 10 cycles as a function of frequency. Decibel baseline-normalized alpha (8 – 12 Hz) power was averaged across all SAM presentations and trials from a pre-defined set of 10 occipital electrodes (excluding electrodes from the midline). We used a 250 ms epoch before the onset of the stimulation in each trial as baseline (−500 ms to −250 ms preceding stimulation onset). Again, for each subject we divided trials in two equally-sized groups (low and high power). The number of perceptual switches was then determined in both groups and compared using a paired-samples t-test.

To determine electrophysiological responses to our stimulation conditions, for each participant we identified the electrodes over visual areas that revealed the strongest alpha and gamma steady-state visually evoked potentials (SSVEPs). Data derived from 10 and 40 Hz stimulation were epoched and analyzed separately. Decibel baseline-normalized (see Cohen, 2014) alpha (9.5 – 10.5 Hz) and gamma (39.5 – 40.5 Hz) power was calculated for each electrode, averaged across all SAM presentations and trials. We used 500 ms time-windows preceding each SAM as the stimulation signal and 250 ms before onset of the stimulation in each trial as baseline (−500 ms to −250 ms before stimulation onset). From a pre-defined set of 10 occipital electrodes we determined for each subject the electrodes that revealed strongest power changes to alpha (10 Hz) and gamma (40 Hz) stimulation. Because we were interested in gamma connectivity between left and right hemispheres, the electrode that mirrored the strongest gamma electrode on the other hemisphere was selected as second gamma electrode.

The extent of spectral coherence in the stimulation conditions was determined for the two selected gamma electrodes. For this, we calculated spectral coherence between the two electrodes in a −500 ms to +1000 ms time-window relative to the onset of each SAM. Coherence values measured during the control condition were then subtracted from coherence values measured during the stimulation conditions (Δ gamma connectivity).

Alpha power changes were calculated for each condition as described above by extracting time-series EEG data from the best alpha electrode. Morlet wavelets spanned from 6 – 14 Hz in 20 steps.

## Results

### Gamma connectivity predicts direction of perceived motion

We analyzed control condition data (no sensory entrainment) in order to replicate previous findings regarding gamma- and alpha-band oscillations associated with visual perception. In line with previous findings, interhemispheric connectivity in the gamma band was related to the direction of perceived motion in SAM. A median-split of the data and a subsequent paired-samples t-test revealed that trials with low gamma connectivity (coherence: M = 0.49, SD = 0.19) are more often perceived as vertical motion, compared to trials with high gamma connectivity (coherence: M = 0.65, SD = 0.14; t(28) = 3.008, p = 0.006; Figure 2 A). Perception of horizontal motion, which crosses the retinal midline, is thought to involve a stronger connection between visual areas in the two hemispheres. Our finding confirms this assumption and presents validation of the chosen study design.

**Figure 2.**
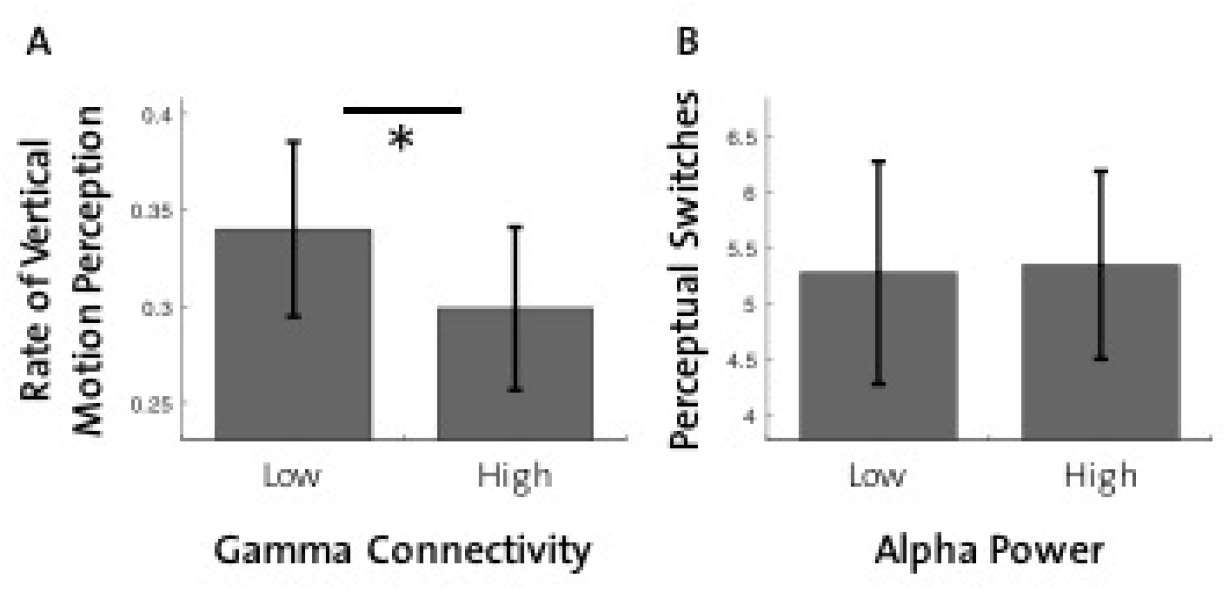
Behavioral correlates of gamma connectivity and alpha power in the control condition. A: More vertical motion of SAM is perceived in trials with low gamma connectivity compared to high gamma connectivity. B: Alpha power level is not predictive for the number of perceptual switches in our study design. (* = p < 0.01, error bars = SEM)

On the other hand, the number of perceptual switches did not differ between low (power: M = 4.41, SD = 4.01) and high (power: M = 10.07, SD = 4.32) alpha power trials as was hypothesized by previous findings (t(28) = −0.120, p = 0.905; Figure 2 B).

### Rhythmic visual stimulation increases alpha power but has no effect on gamma connectivity

In order to determine whether our 10 and 40 Hz stimulation conditions lead to the hypothesized changes in alpha power and gamma connectivity, we first inspect topographical plots and power spectra derived from time-series EEG data during stimulation. The highest power values were found over lateral occipital areas for both stimulation conditions.

In Figure 3 gamma connectivity and alpha power changes across all stimulation conditions (in comparison to control) are depicted. Statistical analysis revealed no effect of visual flicker on interhemispheric gamma connectivity (Δ gamma connectivity; F(3,25) = 0.532, p = 0.665). On the other hand, alpha power was significantly different across stimulation conditions (Δ alpha power; F(3,25) = 9.303, p < 0.001). Bonferroni-corrected pairwise comparisons revealed 10 Hz rhythmic stimulation resulted in significantly higher Δ alpha power compared to all other stimulation conditions (10 Hz vs. 10 Hz jittered: p < 0.001; 10 Hz vs. 40 Hz: p = 0.001; 10 Hz vs. 40 Hz jittered: p = 0.031).

**Figure 3.**
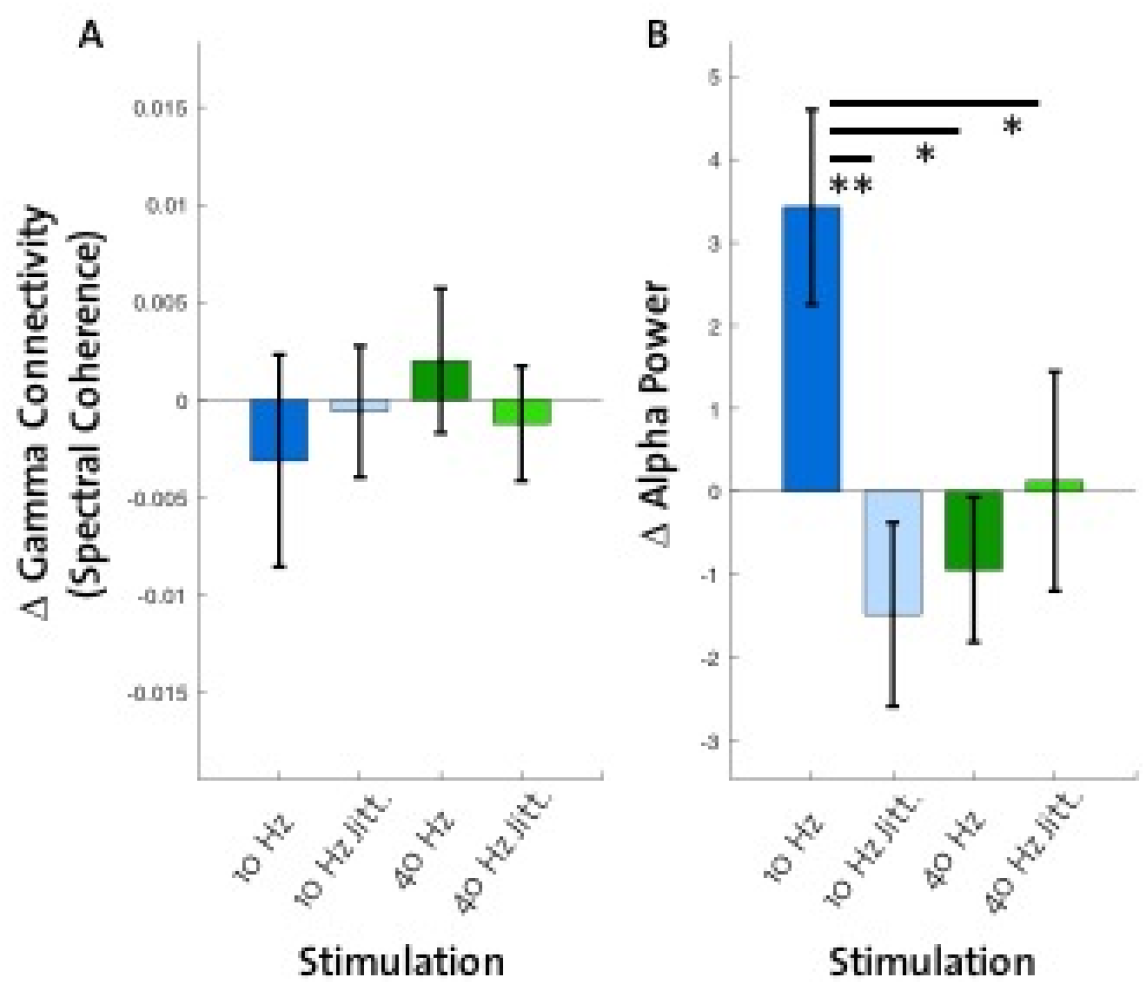
Effect of all stimulation conditions on gamma connectivity and alpha power. A: Gamma connectivity changes compared to control reveal no effect of stimulation. B: 10 Hz stimulation leads to strongest increase (compared to control) in alpha power across stimulation conditions. (** = p < 0.001, * = p < 0.01, error bars = SEM)

### No effect of rhythmic visual stimulation on perceived motion direction or perceptual switches

Contrary to our hypotheses, rhythmic visual stimulation had no effect on perception across all conditions. As we found no effect on Δ gamma connectivity, it may come as no surprise that the stimulation conditions did not differ with respect to the rate of vertical motion perception (F(3,26) = 0.614, p = 0.612; Figure 4 A) and that there was no effect of stimulation on the number of perceptual switches (F(3,26) = 1.274, p = 0.304; Figure 4 B). In an additional analysis, we investigated the influence of the visual flicker on the above describe brain-behavior relationship, i.e. that low interhemispheric gamma coherence favors the perception of vertical motion. For this, we included data from all stimulation conditions into the analysis and divided trials into two equally sized groups of low- and high-gamma connectivity trials. A paired-samples t-test revealed that gamma coherence did not have an effect on direction of perceived motion during visual stimulation (t(28) = 0.460, p = 0.649).

**Figure 4.**
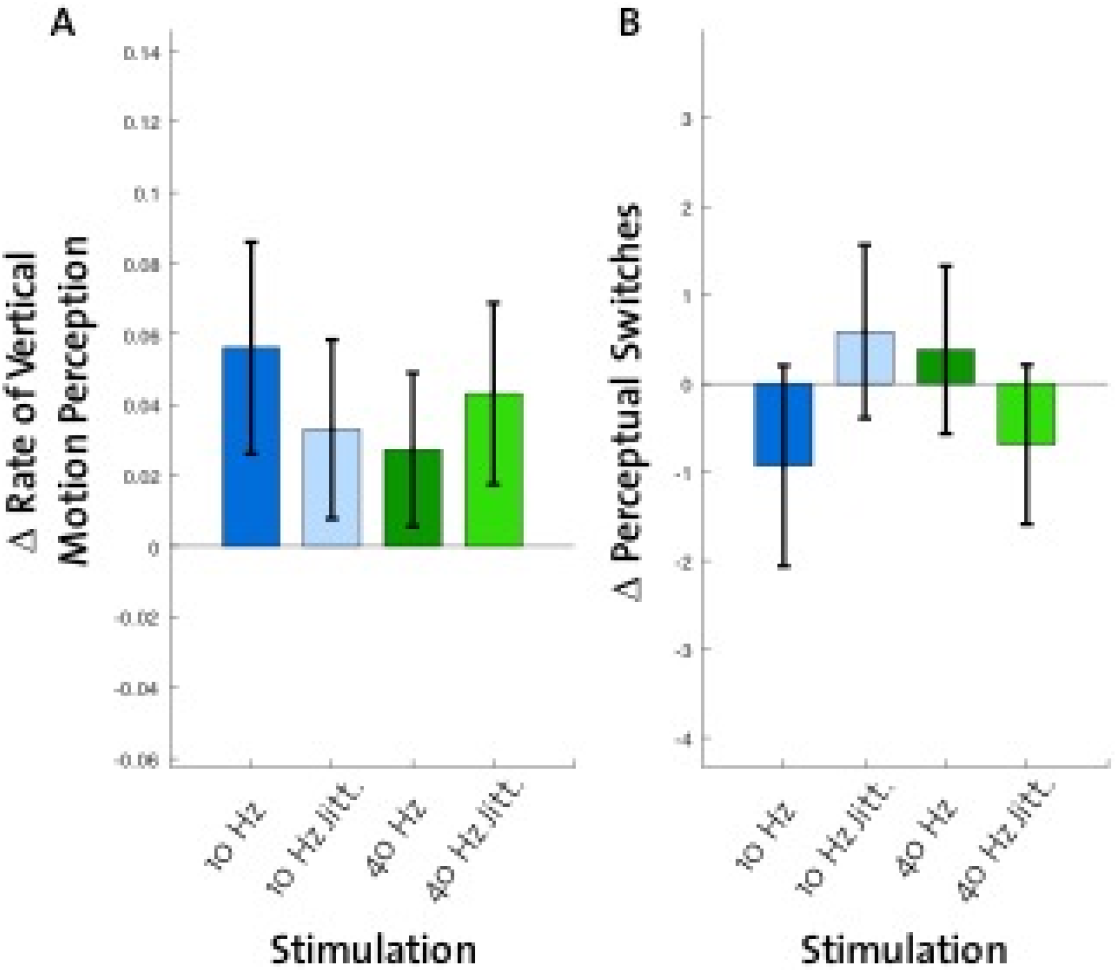
No effect of stimulation conditions on perception. A: Rate of vertical motion perception (compared to control) revealed no differences across stimulation conditions. B: No effect of stimulation was found on changes in perceptual switches compared to control. (error bars = SEM)

## Discussion

In the present study we investigated the effects of bilateral visual stimulation on neuronal activity in the visual system and its modulation of subjective perception of bistable motion. Based on previous literature, we hypothesized a causal role of interhemispheric gamma connectivity on feature integration across both hemispheres (Engel, König, Kreiter, & Singer, 1991), thus being predictive of the motion direction of a bistable apparent motion. Further, we expected occipital alpha power increases to reduce the number of subjective direction reversals of apparent motion. To this end, we entrained brain oscillations in the visual system by presenting flickering LEDs in the gamma (40 Hz) and alpha (10 Hz) frequency band bilaterally while observers reported whether they perceived the motion direction of a stroboscopic alternative motion (SAM) as horizontal or vertical.

### Gamma connectivity predicts direction of perceived motion

In our paradigm, the switch between horizontal and vertical apparent motion is likely to involve a change in interhemispheric connectivity, because information from both hemispheres has to be integrated in the case of perceived horizontal motion (Fries, 2009). As demonstrated previously in a correlative EEG study (Rose & Büchel, 2005), interhemispheric gamma band coherence is increased during perceived horizontal motion compared to vertical motion in SAM, with a peak at 40 Hz. To replicate this finding, we measured phase-based connectivity (spectral coherence) around 40 Hz (36 – 44 Hz) in our control condition and compared perception in low versus high coherence trials. In line with our hypothesis, high gamma connectivity was associated with more horizontal motion perception compared to low gamma connectivity. This finding provides further evidence that gamma oscillations represent a mechanism to facilitate integration of distributed neuronal ensembles enabling behavioral functions such as visual attention and perception (Fries, 2009; Singer, 1999; Tallon-Baudry, 2009). However, this effect is disrupted if the SAM is observed *while* flickering stimuli are presented, irrespective of the flickering frequency. This suggests that visual stimulation itself evokes gamma activity which is unrelated to processing the SMA stimuli and is likely to mask associations between the gamma coherence level and the participant’s percept (Singer, 1999). Moreover, previous literature has found gamma to be susceptible to attentional modulation (e.g. Kohler et al., 2008), which could have occurred in our study because the flickering in the visual periphery might have acted as a distractor.

### Study design does not promote direct perceptual switches

Apart from gamma activity, alpha-band power has also been repeatedly found to correlate (negatively) with perceptual changes of bistable patterns (Başar-Eroglu, Strüber, Kruse, Başar, & Stadler, 1996; Isoglu-Alkaç et al., 2000; Piantoni et al., 2017; Strüber & Herrmann, 2002). However, our control condition did not reveal a difference in the number of perceptual switches preceded by low and high alpha power. A similar absence of the alpha-power-effect was reported by Cabral-Calderin et al. (2015). The authors argue that alpha power might not be causally relevant for perceptual switches, but rather linked to processes of decision-making, motor preparation, or attention (Cabral-Calderin et al., 2015). More research is needed to clarify the underlying role(s) of alpha oscillations in bistable motion perception. Regarding our results, we have to stress that the chosen study design differed in a crucial way from other studies that reported alpha-power effects on perceptual switches. We adapted our design from an fMRI study that investigated neural connections contributing to interhemispheric integration (Genc et al., 2011) and presented SAM with various horizontal-vertical ratios, each for a short time period, and each followed by a decision screen. By contrast findings confirming alpha power changes during perceptual switches come from studies using a paradigm that had participants to observe a bistable motion (e.g. SAM) or ambiguous figure (e.g. Necker cube) until a reversal in perception occurs (Isoglu-Alkaç et al., 2000; Piantoni et al., 2017; Strüber et al., 2014; VanRullen, Reddy, & Koch, 2006). This offers more time to shift into a relaxed state, which is known to be associated with increased alpha-band activity (Adrian & Yamagiwa, 1935). It is possible that this increase is necessary to exert a causal influence on bistable perception.

### Rhythmic sensory stimulation reveals resonance-like response in visual system

In order to test the causal role of alpha and gamma oscillations on bistable motion perception, we aimed to entrain neural activity with sensory rhythmic stimulation. Previous studies using tACS have shown that tACS-induced desynchronization of gamma oscillations (40 Hz) between hemispheres biases the perception of SAM toward vertical motion (Strüber et al., 2014). Helfrich et al. (2014) used 40 Hz tACS in- or out-of-phase across hemispheres and found that these stimulation conditions induced opposite effects on the perception of SAM. While in-phase stimulation increased horizontal motion perception, anti-phase tACS led to a decrease (Helfrich, Knepper, et al., 2014). Although it has been shown that rhythmic sensory stimulation depends on ongoing oscillations (Notbohm et al., 2016; Wälti, Bächinger, Ruddy, et al., 2019), thus representing entrainment of rhythmic neural activity, and that sensory processes can be modulated with this method (Wälti, Bächinger, & Wenderoth, 2019), to our knowledge, no study has used sensory entrainment in a bistable perception paradigm yet.

While rhythmic 10 Hz stimulation revealed the hypothesized increase in alpha power, bilateral rhythmic 40 Hz stimulation had no effect on interhemispheric gamma connectivity, therefore, it comes as no surprise that our stimulation condition did not have an effect on direction of apparent motion or number of perceptual switches. One explanation for the lack of a 40 Hz stimulation effect on interhemispheric coherence might stem from the resonance mechanism in neural populations in the visual system. Studies using steady-state sensory stimulation have shown that each sensory system responds maximally to a specific stimulation frequency (for a review, see Vialatte et al., 2010). In the visual system, alpha activity (around 10 Hz) is thought to represent a mechanism for optimally processing sensory information (Hutcheon & Yarom, 2000; Regan, 1989). In addition, individual stimulation frequencies within a frequency band are thought to be crucial for entrainment effects. This effect has been shown for the individual alpha frequency (IAF) in the visual system (Gulbinaite, van Viegen, Wieling, Cohen, & VanRullen, 2017; Notbohm et al., 2016) and for the individual beta frequency (IBF) in the somatosensory system (Wälti, Bächinger, Ruddy, et al., in 2019; Wälti, Bächinger, & Wenderoth, 2019). In both systems, stimulation frequencies closer to individual ongoing brain oscillations result in stronger entrainment effects and are more likely to modulate perceptual and attentional processes. We therefore argue that future studies using rhythmic sensory stimulation as an entrainment method to modulate perceptual processes should limit the range of stimulation to the system’s resonance frequency band and provide individually adjusted frequencies based on endogenous neural activity in the targeted sensory system.

### Conclusion

Our study provides further evidence that interhemispheric gamma connectivity in the visual system has a crucial role in featuring integration across both hemispheres. However, we also show that it is very difficult to evoke behavioral entrainment effects using rhythmic sensory stimulation.

## Author Contributions

MJW and MB designed research; MJW performed research and analyzed data; MJW, MB, and NW wrote the paper.

## Acknowledgments

We thank Alexandra Bürgler, Lena Salzmann, and Alexander Hess for their help with data acquisition. We also acknowledge the support of the Neuroscience Center Zurich (ZNZ).

## References

Adrian, E. D., & Yamagiwa, K. (1935). The origin of the Berger rhythm. Brain: A Journal of Neurology, 58(3), 323–351.

Başar-Eroglu, C., Strüber, D., Kruse, P., Başar, E., & Stadler, M. (1996). Frontal gamma-band enhancement during multistable visual perception. International Journal of Psychophysiology, 24(1-2), 113–125.

Bressler, S. L., Coppola, R., & Nakamura, R. (1993). Episodic multiregional cortical coherence at multiple frequencies during visual task performance. Nature, 366(6451), 153–156.

Cabral-Calderin, Y., Schmidt-Samoa, C., & Wilke, M. (2015). Rhythmic gamma stimulation affects bistable perception. Journal of Cognitive Neuroscience, 27(7), 1298–1307. doi:10.1162/jocn_a_00781

Capilla, A., Pazo-Alvarez, P., Darriba, A., Campo, P., & Gross, J. (2011). Steady-state visual evoked potentials can be explained by temporal superposition of transient event-related responses. PLoS One, 6(1), e14543. doi:10.1371/journal.pone.0014543

Chaudhuri, A., & Glaser, D. A. (2009). Metastable motion anisotropy. Visual Neuroscience, 7(5), 397–407. doi: 10.1017/s0952523800009706

Cohen, M. X. (2014). Analyzing neural time series data: theory and practice: MIT press.

Doesburg, S. M., Kitajo, K., & Ward, L. M. (2005). Increased gamma-band synchrony precedes switching of conscious perceptual objects in binocular rivalry. Neuroreport, 16(11), 1139–1142.

Engel, A. K., Fries, P., & Singer, W. (2001). Dynamic predictions: oscillations and synchrony in top-down processing. Nature Reviews Neuroscience, 2(10), 704.

Engel, A. K., König, P., Kreiter, A. K., & Singer, W. (1991). Interhemispheric synchronization of oscillatory neuronal responses in cat visual cortex. Science, 252(5009), 1177–1179.

Flevaris, A. V., Martinez, A., & Hillyard, S. A. (2013). Neural substrates of perceptual integration during bistable object perception. Journal of Vision, 13(13), 17. doi:10.1167/13.13.17

Foxe, J. J., & Snyder, A. C. (2011). The Role of Alpha-Band Brain Oscillations as a Sensory Suppression Mechanism during Selective Attention. Frontiers in Psychology, 2, 154. doi:10.3389/fpsyg.2011.00154

Fries, P. (2009). Neuronal gamma-band synchronization as a fundamental process in cortical computation. Annual Review of Neuroscience, 32, 209–224. doi:10.1146/annurev.neuro.051508.135603

Gazzaniga, M. S. (2000). Cerebral specialization and interhemispheric communication: does the corpus callosum enable the human condition? Brain, 123(7), 1293–1326.

Genc, E., Bergmann, J., Singer, W., & Kohler, A. (2011). Interhemispheric connections shape subjective experience of bistable motion. Current Biology, 21(17), 1494–1499. doi:10.1016/j.cub.2011.08.003

Gulbinaite, R., van Viegen, T., Wieling, M., Cohen, M. X., & VanRullen, R. (2017). Individual Alpha Peak Frequency Predicts 10 Hz Flicker Effects on Selective Attention. Journal of Neuroscience, 37(42), 10173–10184. doi:10.1523/JNEUROSCI.1163-17.2017

Haegens, S., & Zion Golumbic, E. (2018). Rhythmic facilitation of sensory processing: A critical review. Neuroscience & Behavioral Reviews, 86, 150–165. doi: 10.1016/j.neubiorev.2017.12.002

Helfrich, R. F., Knepper, H., Nolte, G., Sengelmann, M., Konig, P., Schneider, T. R., & Engel, A. K. (2016). Spectral fingerprints of large-scale cortical dynamics during ambiguous motion perception. Human Brain Mapping, 37(11), 4099–4111. doi:10.1002/hbm.23298

Helfrich, R. F., Knepper, H., Nolte, G., Struber, D., Rach, S., Herrmann, C. S.,… Engel, A. K. (2014). Selective modulation of interhemispheric functional connectivity by HD-tACS shapes perception. PLoS Biol, 12(12), e1002031. doi:10.1371/journal.pbio.1002031

Helfrich, R. F., Schneider, T. R., Rach, S., Trautmann-Lengsfeld, S. A., Engel, A. K., & Herrmann, C. S. (2014). Entrainment of brain oscillations by transcranial alternating current stimulation. Current Biology, 24(3), 333–339. doi:10.1016/j.cub.2013.12.041

Herrmann, C. S., Rach, S., Neuling, T., & Strüber, D. (2013). Transcranial alternating current stimulation: a review of the underlying mechanisms and modulation of cognitive processes. Frontiers in Human Neuroscience, 7, 279. doi:10.3389/fnhum.2013.00279

Hipp, J. F., Engel, A. K., & Siegel, M. (2011). Oscillatory synchronization in large-scale cortical networks predicts perception. Neuron, 69(2), 387–396. doi:10.1016/j.neuron.2010.12.027

Hutcheon, B., & Yarom, Y. (2000). Resonance, oscillation and the intrinsic frequency preferences of neurons. Trends in Neuroscience, 23, 216–222.

Isoglu-Alkaç, Ü., Basar-Eroglu, C., Ademoglu, A., Demiralp, T., Miener, M., & Stadler, M. (2000). Alpha activity decreases during the perception of Necker cube reversals: an application of wavelet transform. Biological cybernetics, 82(4), 313–320.

Isoglu-Alkac, U., & Struber, D. (2006). Necker cube reversals during long-term EEG recordings: subbands of alpha activity. International Journal of Psychophysiology, 59(2), 179–189. doi:10.1016/j.ijpsycho.2005.05.002

Jensen, O., & Mazaheri, A. (2010). Shaping functional architecture by oscillatory alpha activity: gating by inhibition. Frontiers in Human Neuroscience, 4, 186. doi:10.3389/fnhum.2010.00186

Keitel, C., Quigley, C., & Ruhnau, P. (2014). Stimulus-driven brain oscillations in the alpha range: entrainment of intrinsic rhythms or frequency-following response? Journal of Neuroscience, 34(31), 10137–10140. doi:10.1523/JNEUROSCI.1904-14.2014

Kohler, A., Haddad, L., Singer, W., & Muckli, L. (2008). Deciding what to see: the role of intention and attention in the perception of apparent motion. Vision Research, 48(8), 1096–1106. doi: 10.1016/j.visres.2007.11.020

Lange, J., Keil, J., Schnitzler, A., van Dijk, H., & Weisz, N. (2014). The role of alpha oscillations for illusory perception. Behavioural Brain Research, 271, 294–301. doi:10.1016/j.bbr.2014.06.015

Liu, Q., Ganzetti, M., Wenderoth, N., & Mantini, D. (2018). Detecting Large-Scale Brain Networks Using EEG: Impact of Electrode Density, Head Modeling and Source Localization. Frontiers in Neuroinformatics, 12, 4. doi:10.3389/fninf.2018.00004

Mima, T., Oluwatimilehin, T., Hiraoka, T., & Hallett, M. (2001). Transient interhemispheric neuronal synchrony correlates with object recognition. Journal of Neuroscience, 21(11), 3942–3948.

Notbohm, A., Kurths, J., & Herrmann, C. S. (2016). Modification of Brain Oscillations via Rhythmic Light Stimulation Provides Evidence for Entrainment but Not for Superposition of Event-Related Responses. Frontiers in Human Neuroscience, 10, 10. doi:10.3389/fnhum.2016.00010

Piantoni, G., Romeijn, N., Gomez-Herrero, G., Van Der Werf, Y. D., & Van Someren, E. J. W. (2017). Alpha Power Predicts Persistence of Bistable Perception. Scientific Reports, 7(1), 5208. doi:10.1038/s41598-017-05610-8

Ramachandran, V. S., & Anstis, S. M. (1983). Perceptual organization in moving patterns. Nature, 304(5926).

Regan, D. (1977). Steady-state evoked potentials. Journal of Optical Society of America, 67(11), 1475–1489.

Regan, D. (1989). Human brain electrophysiology: evoked potentials and evoked magnetic fields in science and medicine. New York: Elsevier.

Rose, M., & Büchel, C. (2005). Neural coupling binds visual tokens to moving stimuli. Journal of Neuroscience, 25(44), 10101–10104. doi:10.1523/JNEUROSCI.2998-05.2005

Sangiuliano Intra, F., Avramiea, A. E., Irrmischer, M., Poil, S. S., Mansvelder, H. D., & Linkenkaer-Hansen, K. (2018). Long-Range Temporal Correlations in Alpha Oscillations Stabilize Perception of Ambiguous Visual Stimuli. Frontiers in Human Neuroscience, 12, 159. doi:10.3389/fnhum.2018.00159

Schwab, K., Ligges, C., Jungmann, T., Hilgenfeld, B., Haueisen, J., & Witte, H. (2006). Alpha entrainment in human electroencephalogram and magnetoencephalogram recordings. Neuroreport, 17(17), 1829–1833.

Singer, W. (1999). Neuronal synchrony: a versatile code for the definition of relations? Neuron, 24(1), 49–65.

Srinivasan, R., Russell, D. P., Edelman, G. M., & Tononi, G. (1999). Increased synchronization of neuromagnetic responses during conscious perception. Journal of Neuroscience, 19(13), 5435–5448.

Strüber, D., Başar-Eroglu, C., Hoff, E., & Stadler, M. (2000). Reversal-rate dependent differences in the EEG gamma-band during multistable visual perception. International Journal of Psychophysiology, 38(3), 243–252.

Strüber, D., & Herrmann, C. S. (2002). MEG alpha activity decrease reflects destabilization of multistable percepts. Cognitive Brain Research, 14(3), 370–382.

Strüber, D., Rach, S., Trautmann-Lengsfeld, S. A., Engel, A. K., & Herrmann, C. S. (2014). Antiphasic 40 Hz oscillatory current stimulation affects bistable motion perception. Brain Topography, 27(1), 158–171. doi:10.1007/s10548-013-0294-x

Tallon-Baudry, C. (2009). The roles of gamma-band oscillatory synchrony in human visual cognition. Frontiers in Bioscience, 14, 321–332.

Thut, G., Schyns, P. G., & Gross, J. (2011). Entrainment of perceptually relevant brain oscillations by non-invasive rhythmic stimulation of the human brain. Frontiers in Psychology, 2, 170. doi:10.3389/fpsyg.2011.00170

VanRullen, R., Reddy, L., & Koch, C. (2006). The continuous wagon wheel illusion is associated with changes in electroencephalogram power at approximately 13 Hz. Journal of Neuroscience, 26(2), 502–507. doi:10.1523/JNEUROSCI.4654-05.2006

Vialatte, F. B., Maurice, M., Dauwels, J., & Cichocki, A. (2010). Steady-state visually evoked potentials: focus on essential paradigms and future perspectives. Progress in Neurobiology, 90(4), 418–438. doi:10.1016/j.pneurobio.2009.11.005

Wälti, M. J., Bächinger, M., Ruddy, K. L., & Wenderoth, N. (2019). Steady-state responses in the somatosensory system interact with endogenous beta activity. bioRxiv 690495. doi:10.1101/690495

Wälti, M. J., Baechinger, M., & Wenderoth, N. (2019). Modulation of tactile detection threshold with rhythmic somatosensory entrainment. bioRxiv, 695692

Zoefel, B., Ten Oever, S., & Sack, A. T. (2018). The Involvement of Endogenous Neural Oscillations in the Processing of Rhythmic Input: More Than a Regular Repetition of Evoked Neural Responses. Frontiers in Neuroscience, 12, 95. doi:10.3389/fnins.2018.00095

